# Regional susceptibility of PV interneurons in an hAPP-KI mouse model of Alzheimer’s disease pathology

**DOI:** 10.1101/2025.04.01.646485

**Authors:** Mercedes M. Gonzalez, Benjamin Magondu, Matthew J. M. Rowan, Craig R. Forest

## Abstract

Early-stage Alzheimer’s pathology correlates with disrupted neuronal excitability, which can drive network and cognitive dysfunction even prior to neurodegeneration. However, the emergence and extent of these changes may vary by brain region and cell types situated in those regions. Here we aimed to investigate the effects of AD pathology on different neuron subtypes in both the entorhinal cortex, a region with enhanced pathology in early AD, and the primary visual cortex, a relatively unaffected region in early-stage AD. We designed and employed a semi-automated patch clamp electrophysiology apparatus to record from fast-spiking parvalbumin interneurons and excitatory neurons in these regions, recording from over 150 cells in young adult APP-KI mice. In entorhinal cortex, amyloid overproduction resulted in PV interneuron hypoexcitability, whereas excitatory neurons were concurrently hyperexcitable. Conversely, neurons of either subclass were largely unaffected in the visual cortex. Together, these findings suggest that fast-spiking parvalbumin interneurons in the entorhinal cortex, but not in the visual cortex, play an integral role in AD progression.

## Introduction

Alzheimer’s disease (AD) is a devastating neurodegenerative condition characterized by progressive cognitive decline and memory loss. As the leading cause of dementia, AD poses significant challenges for healthcare systems globally, necessitating urgent attention and innovative approaches to diagnosis, treatment, and care ***Breijyeh and Karaman (2020a); Hurd et al. (2013***). AD is hallmarked by the accumulation of amyloid-beta in the disease early-stages, followed by the emergence of intraneuronal tau tangles later on which are thought to be casually linked neuronal damage ***Breijyeh and Karaman (2020a***,b); ***Belloy et al. (2019); Knopman et al. (2021***). While this hypothesis has provided valuable insights into the molecular underpinnings of AD and guided much of the drug development efforts in the field, it has also faced significant challenges and limitations in the clinic ***Sims et al. (2023); Van Dyck et al. (2023); Kim et al. (2022***). Despite extensive research and clinical trials targeting amyloid and tau pathology, effective disease-modifying therapies are still lacking, likely owing to the complex pathophysiology of AD. A major reason why advancement has been slow may be due to the vast diversity of brain cell types, wherein each type may be selectively vulnerable to different forms of AD pathology.

The emergence of hAPP/Abeta pathology is generally regarded as the first factor in the AD cascade, in both the familial and sporadic forms of the disease ***Selkoe and Hardy (2016***). In recent years, there has been growing appreciation of the multimodal effects of early AD pathogenesis. This includes correlations between increased Abeta and aberrant circuit function, as well as neuroinflammation, impaired clearance mechanisms, among other outcomes ***Knopman et al. (2021); Cubinkova et al. (2018***). Relevant to our study here, evidence from both animal models and clinical studies provides compelling support for the involvement of neuronal circuit dysregulation in AD ***Busche et al. (2008, 2015***). Several clinical studies indicate that there is increased brain activity in both sporadic as well as familial AD ***Celone et al. (2006); Bassett et al. (2006***). Early Abeta pathology results in neuronal and synaptic impairment ***Busche et al. (2012***), leading to deficits in memory and executive function ***Vossel et al. (2016); Sanchez et al. (2012); Vossel et al. (2021***). Animal models of AD have also revealed alterations in neuronal network activity preceding the onset of classical pathological hallmarks ***Busche et al. (2015)***; ***Shimojo et al. (2020***). Hyperexcitability in AD has been hypothesized to result from disrupted balance in network excitatory and inhibitory activity, through various cellular and synaptic mechanisms ***Bai et al. (2017***); ***Targa Dias Anastacio et al. (2022***). In the context of AD, disruptions in the function of GABAergic interneurons are proposed to contribute to circuit hyperexcitability and cognitive impairment ***Arroyo-Garcia et al. (2021)***; ***Nuriel et al. (2017)***; ***Li et al. (2021)***; ***Sederberg et al. (2007***).

While region-specific vulnerabilities in AD have been researched over the past several decades ***Braak and Braak (1991); Mrdjen et al. (2019***), it remains unclear why some brain regions are more severely affected by AD. The lateral entorhinal cortex (LEC) is the first cortical brain region to exhibit AD pathology ***Khan and et al. (2014)***; ***Kobro-Flatmoen et al. (2016***). There has been increasing evidence of vulnerability in the LEC ***Karimani et al. (2024***); there is a marked reduction in Layer II neurons in mild AD ***Braak and Del Tredici (2012); Gómez-Isla et al. (1996***), Abeta and tau accumulate in the entorhinal cortex early in disease progression ***Thal et al. (2002); Harrison et al. (2019***), and local LEC circuits are hyperexcitable in early stage AD ***Duffy et al. (2015***).

Understanding the regional and temporal emergence of interneuron dysfunction is crucial for deciphering the complex pathophysiology of AD and identifying potential therapeutic targets. More-over, temporal changes in interneuron function and connectivity dynamics may underlie the progressive nature of cognitive decline in AD, with dysfunction manifesting at different disease stages. Here we investigated the regional susceptibility of interneuron and excitatory neurons to early Abeta pathology in distinct AD-vulnerable and control regions, by quantifying both the passive and active intrinsic properties of PV interneurons and excitatory neurons at an early disease time point. This study not only enhances our fundamental understanding of AD progression, but also guides future development of more effective diagnostic and therapeutic strategies.

## Results

The lateral entorhinal cortex (LEC) is one of the first brain regions to exhibit early neurodegeneration and neurofibrillary tangles in AD ***Igarashi (2023)***; ***Gómez-Isla and et al. (1996***); ***Khan and et al. (2014***). Furthermore, recent evidence has suggested there may be altered excitability in this brain region in early AD in humans ***Khan and et al. (2014***) as well as in several mouse models ***Khan and et al. (2014***); ***Nuriel et al. (2017***); ***Goettemoeller et al. (2023***). Using APP-KI mice which produce amyloid pathology (i.e., h*A*beta^**S**we,**A**rc,**A**us^ mice) without disturbing intrinsic APP expression, we recorded from fluorescently-labeled PV interneurons in the EC ***Goettemoeller et al. (2023***) (1A-B) (of 8-10 week old mice) to determine whether this model shows any indication of altered excitability. We found hypoexcitablity of these PV interneurons across nearly all current injections (1C). The passive properties of these cells showed no significant differences (1D). However, the action potential half-width of the first action potential was increased and the maximum dV/dt was decreased (1E) in the SAA mice, together suggesting sodium channel dysfunction, together suggesting changes in active parameters underlie the altered AP firing in LEC PV interneurons.

**Figure 1.**
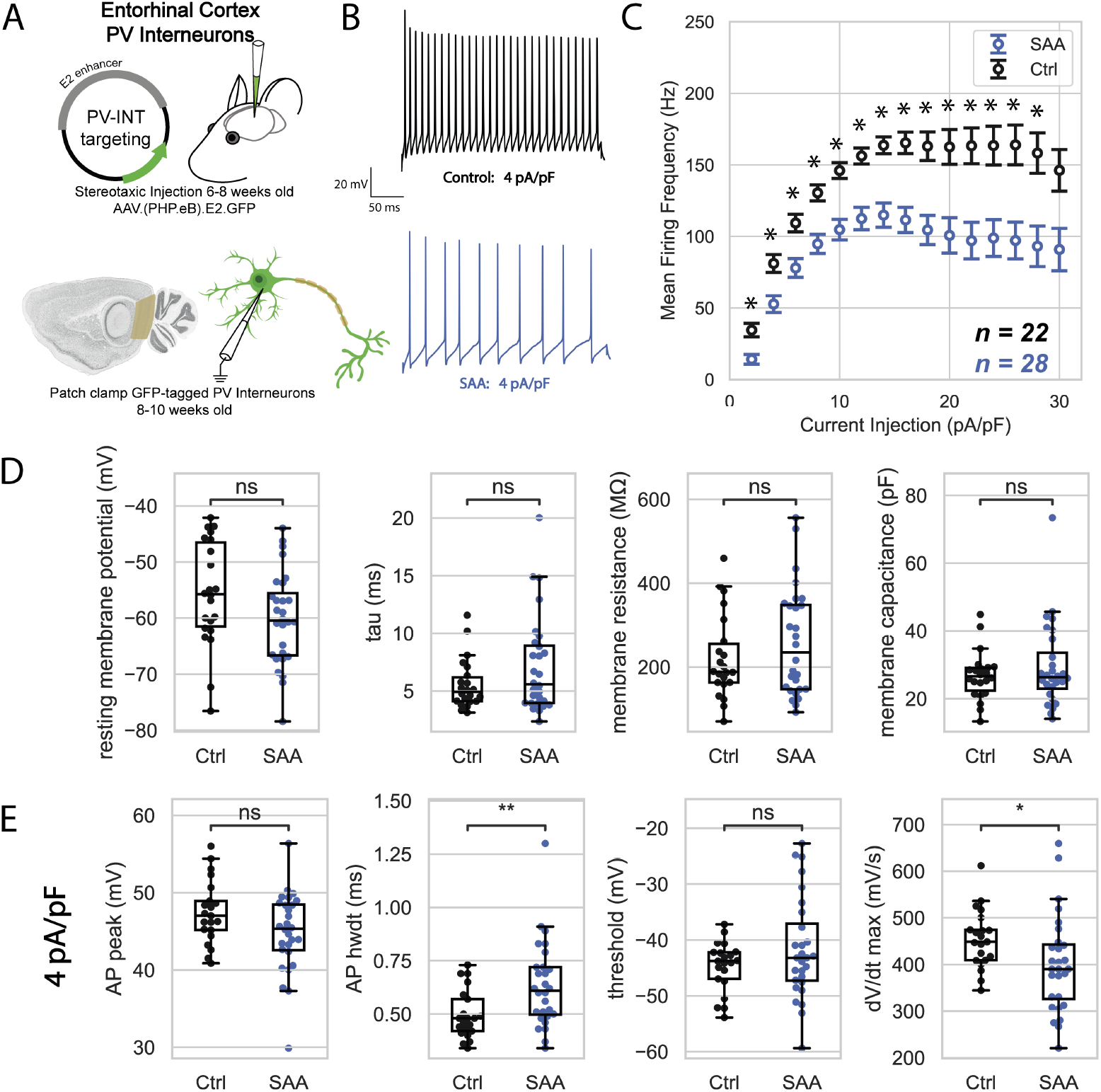
PV Interneurons in the lateral entorhinal cortex exhibit reduced firing. A. Schematic of the experimental approach, including injection of the AAV in the entorhinal cortex and patch clamp of fluorescent PV interneurons. B. Representative current clamp traces from the AD pathology model (SAA, blue) and the corresponding control (Ctrl, black) in response to a 4 pA/pF current injection. C. Summary of mean firing frequency as a function of current injection. SAA exhibits significant hypoexcitability across all current injections (Sidak’s multiple comparison’s test, * indicates p < 0.05). (Ctrl n = 22 cells from 4 mice, SAA n = 28 cells from 3 mice) D. Passive properties of PV interneurons: resting membrane potential, time constant (*τ*), membrane resistance, and membrane capacitance. E. Properties of the first action potential evoked from a 4 pA/pF current injection (from left to right): action potential peak, action potential half-width, threshold, and maximum dV/dt. Action potential half-width is significantly increased in SAA (unpaired t-test p = 0.0073) and maximum dV/dt is decreased (unpaired t-test p = 0.043).

We also considered the intrinsic properties of excitatory neurons across all layers of the lateral entorhinal cortex (2A) to determine if there may be a corresponding compensatory change in excitability. We found that these cells were hyperexcitable near threshold (2B-C) ***Chen et al. (2023***) potentially owing to a number of factors. For example, the membrane capacitance and thus time constant, were both decreased in the SAA mice (2D). These excitatory cells also exhibited shortened action potential half-width (2E) in the SAA mice. This decrease in action potential half-width may be related to altered channel function ***Olah et al. (2021***) at this age in SAA-LEC excitatory neurons.

**Figure 2.**
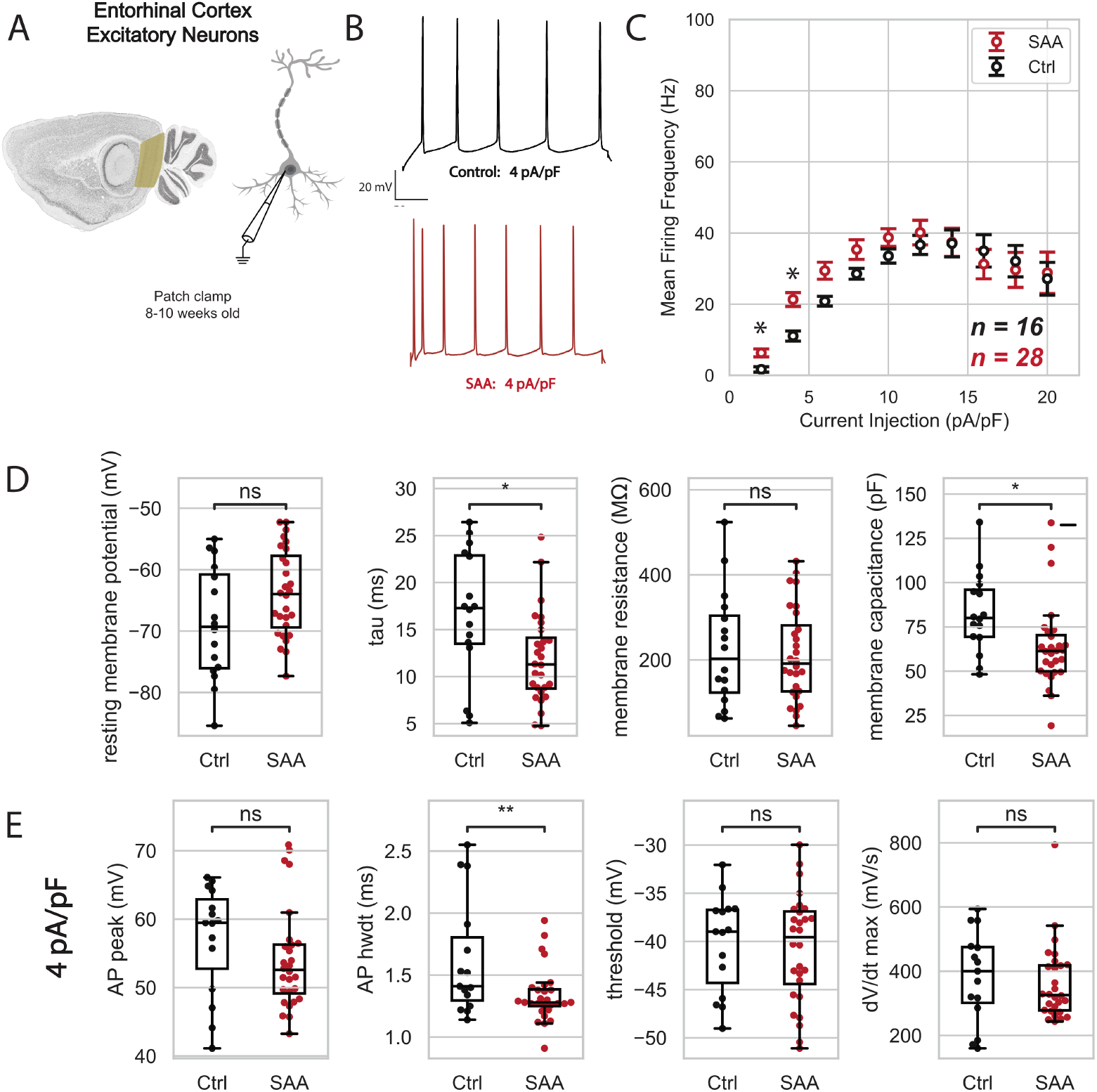
Excitatory neurons in the LEC are hyperexcitable near threshold. A. Schematic of the experimental approach: patch clamp of excitatory neurons. B. Representative current clamp traces from the AD model (SAA, red) and the corresponding control (Ctrl, black) in response to a 4 pA/pF current injection. C. Summary of mean firing frequency as a function of current injection. SAA exhibits slight hyperexcitability near threshold (Sidak’s multiple comparison’s test, * indicates p < 0.05). (Ctrl n = 16 cells from 3 mice, SAA n = 28 cells from 3 mice) D. Passive properties of PV interneurons: resting membrane potential, time constant (τ), membrane resistance, and membrane capacitance. Tau is significantly decreased in SAA (unpaired t-test p = 0.010). Membrane capacitance is decreased in SAA (unpaired t-test p = 0.015). E. Properties of the first action potential evoked from a 4 pA/pF current injection (from left to right): action potential peak, action potential half-width, threshold, and maximum dV/dt. Action potential half-width is significantly decreased in SAA (unpaired t-test p = 0.0094).

To determine whether PV interneuron excitability changes in the SAA line emerged as a function of brain region or rather in a more uniform manner in the cortex, we next recorded from PV interneurons and excitatory neurons across all layers in the primary visual cortex region (3A). In contrast to the LEC, PV interneurons in visual cortex did not show differences in firing rate (3B-C) or passive properties (3D).

**Figure 3.**
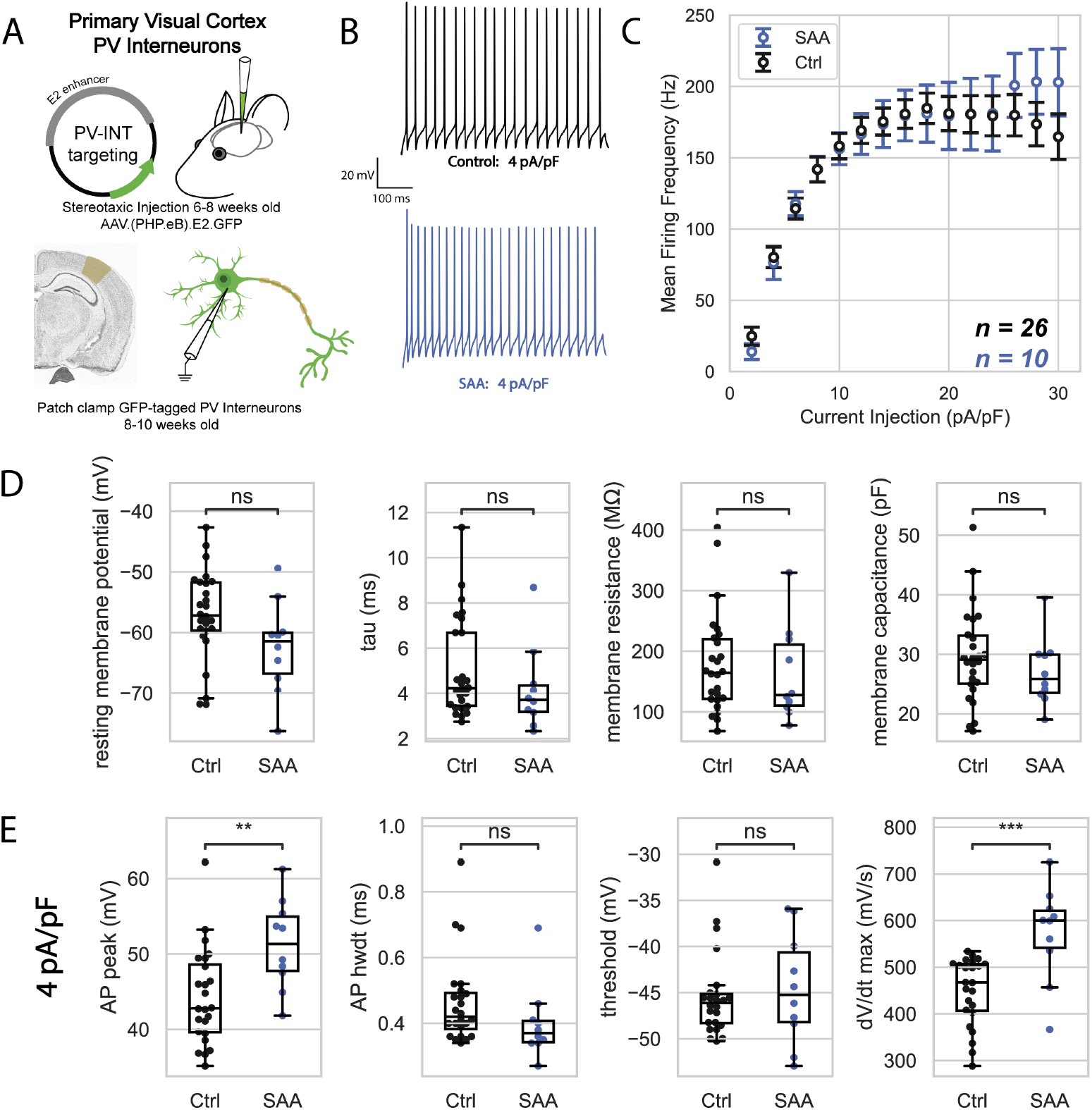
PV interneurons in the primary visual cortex display no differences in firing. A. Schematic of the experimental approach, including injection of the AAV in the primary visual cortex and patch clamp fluorescent PV interneurons. B. Representative current clamp traces from the AD model (SAA, blue) and the corresponding control (Ctrl, black) in response to a 4 pA/pF current injection. C. Summary of mean firing frequency as a function of current injection, with no significant difference in firing frequency. (Ctrl n = 26 cells from 3 mice, SAA n = 10 cells from 3 mice) D. Passive properties of PV interneurons: resting membrane potential, time constant (τ), membrane resistance, and membrane capacitance. E. Properties of the first action potential evoked from a 4 pA/pF current injection (from left to right): action potential peak, action potential half-width, threshold, and maximum dV/dt. Action potential peak (unpaired t-test p = 0.0056) and maximum dV/dt are increased in hAPP (unpaired t-test p = 0.00032)).

Despite the lack of changes in overall intrinsic excitability, AP peak and dV/dt max were both increased in PV interneurons of SAA mice (3E). This increase in AP peak and dV/dt max suggests enhanced sodium channel activity in the primary visual cortex region, potentially reflecting compensatory mechanisms in PV interneurons to maintain circuit stability. This may either represent a homeostatic adaptation to preserve neuronal function in the relatively unaffected region of early Alzheimer’s disease, or local differences in ion channel composition specific to the visual cortex.

Following a similar pattern as in PV cells, the excitatory neurons of the primary visual cortex (4A), showed no differences in intrinsic properties such as membrane capacitance, time constant, or action potential dynamics in the SAA model (4B-C), suggesting that their excitability and passive properties remain largely unaltered in this brain region. However, a reduced resting membrane potential was noted (4D), indicating a hyperpolarized baseline state. This hyperpolarization could reflect subtle shifts in ion channel activity, such as increased potassium channel conductance or altered sodium-potassium pump function, which may serve as a compensatory mechanism to stabilize neuronal activity in a region not typically vulnerable in early AD.

**Figure 4.**
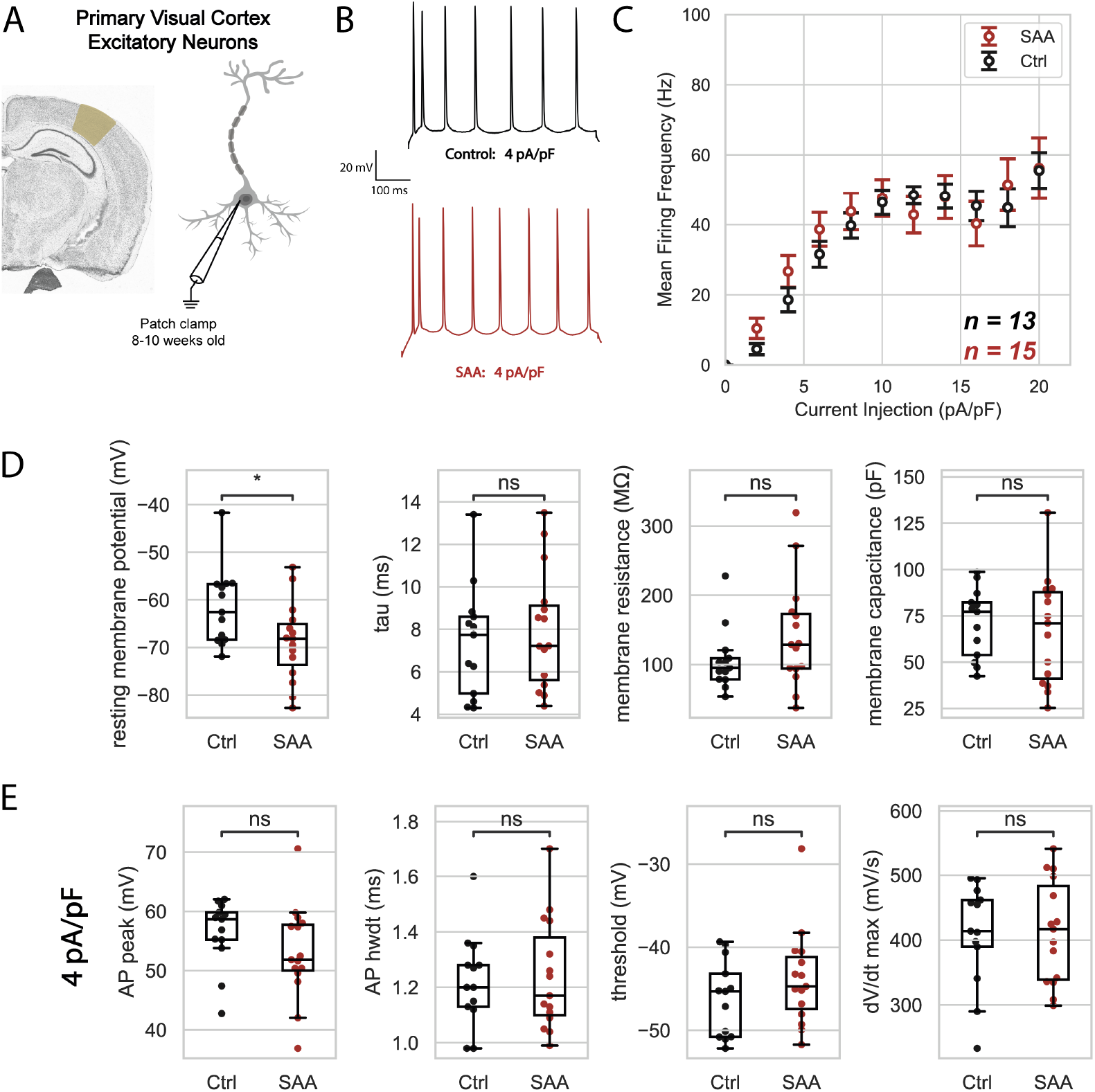
Excitatory neurons in the primary visual cortex are unaffected in SAA mice. A. Schematic of the experimental approach: patch clamp excitatory neurons. B. Representative current clamp traces from the AD model (SAA, red) and the corresponding control (Ctrl, black) in response to a 4 pA/pF current injection. C. Summary of mean firing frequency as a function of current injection, with no differences in firing frequency. (Ctrl n = 13 cells from 2 mice, SAA n = 15 cells from 3 mice) D. Passive properties of excitatory neurons: resting membrane potential, time constant (τ), membrane resistance, and membrane capacitance. Resting membrane potential is slightly decreased in SAA (unpaired t-test p = 0.027). E. Properties of the first action potential evoked from a 4 pA/pF current injection (from left to right): action potential peak, action potential half-width, threshold, and maximum dV/dt.

Together, our results suggest that the excitable properties of neurons in the LEC are more susceptible to change with respect to other cortical areas.

## Discussion

Here we systematically studied the intrinsic properties of parvalbumin interneurons and excitatory neurons across two cortical regions using an hAPP-knock in mouse model of Alzheimer’s disease pathology. We sought to quantify whether early changes in PV interneuron intrinsic excitability in this model ***Petrache et al. (2019***) were differentially expressed by region, thereby acting as potential contributor to the hyperexcitability apparent in early AD pathology ***Busche et al. (2008); Busche and Konnerth (2015); Davis et al. (2014); Giorgio et al. (2024); Roemer-Cassiano et al. (2025***). We found that (at 8-10 weeks of age) PV interneurons do exhibit intrinsic hypoexcitability across the entorhinal cortex but not in the visual cortex. As amyloid levels appear uniform across the entire hippocampal formation and cortex in this model at this early age ***Xia et al. (2022***), this suggests that cell-intrinsic vulnerabilities specific to the entorhinal region underlie our observations.

Our findings support the idea that PV interneurons in the entorhinal cortex are more susceptible to early-stage AD (APP/amyloid) pathology with respect to other cortical areas. Specifically, the increased half-width of the action potentials in SAA mice suggests differences in ion channel function or density, in particular voltage gated sodium and potassium channels ***Olah et al. (2021***); ***Gu et al. (2018)***; ***Hijazi et al. (2020)***; ***Verret et al. (2012)***; ***Martinez-Losa et al. (2018***); ***Manville and Abbott (2021***). The AP peak is driven by the influx of sodium ions through voltage-gated sodium channels. An increase in dV/dt max, the rate of rise of the action potential, suggests that these channels are either opening more rapidly or in greater numbers, resulting in a steeper depolarization phase. These changes could also reflect region-specific compensatory mechanisms in PV interneurons to maintain circuit stability in response to altered network activity or excitatory drive. While active properties were altered in PV cells, we found that PV passive properties were unchanged. This is a distinction from our previous study using adult-onset hAPP expression ***Goettemoeller et al. (2023***). Future work utilizing models to induce amyloid production in mature animals will thus be necessary to examine these mechanistic distinctions. Nonetheless, several recent studies continue to converge on the theme of early cellular vulnerability based on region, in particular, the entorhinal cortex ***Mitchell and Silver (2003); Magee (2000); Taddei and Duff (2025); Mathys et al. (2024***) in AD. We also found a concurrent increase in excitatory neurons excitability near threshold in the entorhinal region, however firing rate was unaffected at higher current injections. Notably, the time constant and membrane capacitance is decreased in this SAA model, suggesting structural changes which may explain the hyperexcitability in these cells. These passive changes, together with the decreased half-width of the action potentials, suggest that both the size of these excitatory neurons and the ion channel activity on these neurons may contribute to their enhanced intrinsic excitability.

Typically, V1 is not commonly studied in early AD since it does not play a major role in memory and cognition. However, V1 is affected in late disease progression, as amyloid and tau are often accumulated across the entire cortex, and visual and motor systems are severely affected in later stage AD ***Liebscher et al. (2016***). Thus we sought to also examine whether PV and excitatory neurons were affected during early pathology in the APP-KI model in the V1 region. In contrast to the entorhinal cortex, our data showed no indication of altered excitability of PV interneurons via firing frequency or passive membrane properties. Similarly, there was no apparent change in the intrinsic excitability of excitatory neurons in V1, aside from a slight difference in the resting membrane potential.

In summary, recent evidence has shown that both PV interneurons ***Olah et al. (2021)***; ***Verret et al. (2012)***; ***Hijazi et al. (2020)***; ***Goettemoeller et al. (2023***) and excitatory neurons ***Chen et al. (2023***) are physiologically vulnerable in AD. This work supports the idea that there is reduced excitability of PV interneurons and concurrent enhancement of excitatory neurons during early AD pathology, and that is phenomenon may preferentially emerge in the entorhinal cortex. The evidence across multiple models of AD supports the overarching hypothesis that PV cells in the EC are particularly vulnerable and interesting early therapeutic targets.

## Methods and Materials

### Alzheimer’s Disease Mouse Model

The hAPP-knock-in mouse model of AD (B6.Cg-Apptm1.1Dnli/J, Jax Strain #034711 ***Xia et al. (2022***)) was used. the B6J hAbeta line (B6.Cg-Appem1Adiuj/J, Jax Strain #033013) was used as a control.

### Stereotaxic viral injections

Mice were injected with AAV(PHP.eB)S5E2.GFP (Addgene #135631) at 6-8 weeks old via streotax (David Kopf Instruments). 300 *µ*L was injected over the duration of 10 minutes + 5 minutes to prevent backflow (Hamilton Neuros Syringes) at the steretaxic coordinates in Table 1. Skin was sutured and mice were monitored during recovery from anesthesia under a heat lamp. Mice received SR buprenorphrine for pain relief. Mice were checked for wound healing and condition for 3 days post operation.

**Table 1.** Stereotaxic coordinates (in mm) by brain region.

| Brain region | A/P | M/L | D/V |
| --- | --- | --- | --- |
| LEC | -4.5 | 3.4 | 2.5 |
| V1 | -2.79 | 2.5 | 1 |

### Acute brain slice preparation

All animal procedures were done in accordance with the US National Institutes of Health Guide for the Care and Use of Laboratory Animals and the Georgia Institute of Technology animal care committee’s regulations. Mice were anesthetized with 4% isoflurane and decapitated. Brain slices (300*µ*m) were prepared (Leica VT1200 Vibratome) in ice cold cutting solution (in mM: 25 NaH CO_3_, 10 Dextrose, 75 Sucrose, 87 NaCl, 2.5 KCl, 1.25 NaH_2_PO4, 1 CaCl_2_, 2 Mg_2_). Slices incubated in 33°C recording solution (in mM: 26.2 NaH CO_3_, 11 Dextrose, 28.6 NaCl, 2.5 KCl, 1 NaH_2_PO_4_, 1.5 CaCl_2_, 1.5 MgCl_2_) for 30 minutes and at 24°C for 15 minutes before the experiment. Solutions were oxygenated with 95% Oxygen/ 5% Carbon Dioxide during the length of the experiment.

### Semi-automated electrophysiology

A standard electrophysiology setup (SliceScope Pro 3000, Scientifica Ltd) with PatchStar micromanipulators at a 24°approach angle was used for all electrophysiology recordings. We used a 40x objective (LUMPFLFL40XW/IR, NA 0.8, Olympus) and Rolera Bolt camera (QImaging), illuminated under DIC with an infrared light-emitting diode (Scientfica). A peristaltic pump (120S/DV, Watson-Marlow) perfused the brain slices with recording solution, as described above. A pressure control box (Neuromatic Devices) regulated internal pipette pressure as well as a custom machined chamber with a smaller side chamber for cleaning solution.

Internal solution (in mM: 138 K-gluconate, 2 KCl, 10 HEPES, 4 Na_2_ATP, 0.5 Na_2_GTP, 4 MgCl_2_, 3 L-Ascorbic Acid) was thawed on ice and filtered (0.2 *µ*m) each day. Borosilicate glass pipettes with internal filament were filled with 5-10 *µ*L of internal solution. Automated cleaning of the pipette between patch attempts was implemented with LabVIEW, as in ***Landry et al. (2020***). 2% w/v Tergazyme solution was used to clean the pipette. The rinsing step was not included. All other steps of the patch clamp process were done manually, with computer-control of the internal pipette pressure.

Cells were injected with current to maintain the membrane potential at −70 mV. For each cell, a voltage clamp membrane test was run to initially estimate the passive properties of the cell. To determine both passive and active properties of the cells, a current clamp protocol with a depolarizing step −20 pA followed by a series of hyperpolarizing 2 pA/pF steps from −2 pA/pF to 30 pA/pF (normalized to the estimated capacitance of the cell) was run.

Recordings with access resistance less than 20 MΩ were considered good quality recordings for this analysis. The average access resistance across all recordings was 14.6 MΩ and the average holding current was −20.0 pA. Pipette capacitance was neutralized and access resistance resistance was compensated via bridge balance for current clamp recordings.

### Statistics and analysis

Python and GraphPad were used to run statistical analyses. Two-tailed unpaired t-tests were used for comparing between groups for both passive and spike parameters. Significant p-values are shown in the figure caption where appropriate. A two-way ANOVA test with Sidak’s multiple comparisons was used to compare the mean firing frequency across current injections. Cells with passive properties with z-scores greater than 3 were considered outliers and excluded from the analysis. Box plots show the mean as the center line, with the box representing the data quartiles (25% and 75%), and the whiskers representing 1.5^*^IQR (interquartile range) away from the quartiles.

## Acknowledgments

NIH Grant 1RF1AG079269-01, NIH BRAIN Initiative Grant (NEI and NIMH 1-U01-MH106027-01), NIH R01NS102727, NIH Single Cell Grant 1 R01 EY023173, and NIH R01DA029639, support from Georgia Tech through the Institute for Bioengineering and Biosciences, Invention Studio, and the George W. Woodruff School of Mechanical Engineering.

